# Polymer-Assisted Condensation: A Mechanism for Hetero-Chromatin Formation and Epigenetic Memory

**DOI:** 10.1101/2021.10.13.464201

**Authors:** Jens-Uwe Sommer, Holger Merlitz, Helmut Schiessel

## Abstract

We consider the formation of droplets from a 2-component liquid mixture induced by a large polymer chain that has preferential solubility with one of the components. We assume that the liquid mixture is in a fully miscible state, but far above the critical interaction limit of the two species. We show that the polymer coil acts as a chemical potential trap, which can shift the mixture inside the polymer volume into the partially miscible state and thus triggers the formation of a polymer-bound droplet of the preferred solvent phase which we denote as polymer-assisted condensation (PAC). We propose a simple mean-field model which can predict the essential feature of PAC and perform molecular-dynamics simulations to show that the predicted phase behavior is robust against fluctuation effects. Our model aims to understand the formation of macromolecular condensates inside the cell nucleus, such as those formed by heterochromatin 1 (HP1). We propose that such droplets organize the spatial structure of chromatin into hetero- and euchromatin and ensure the propagation of epigenetic information through the cell generations.

## I. INTRODUCTION

When a cell divides, not only does the genetic information have to be faithfully duplicated, but its epigenetic information as well has to be passed on to the two daughter cells. Epigenetics is defined as modifications of the functions of genes that are stable and inheritable during mitosis, possibly meiosis (see e.g. [1]). They typically occur in the form of modifications of the DNA itself in the form of DNA methylation and of the histone proteins complexed with the DNA molecules in the form of methylations, acetylations etc. of amino acids [1]. In the following we focus on the latter type of modifications. The histone proteins are associated with the DNA in the form of nucleosomes, DNA-protein complexes where about 147 basepairs (bp) of DNA are wrapped around an octamer of histone proteins [2, 3]. About 3/4 of human DNA is sequestered this way. There exist a number of specific modifications of the histone proteins that assign specific functions to the involved chromosome stretches. For example, the trimethylation of the ninth amino acid (counted from the N-terminus) of histone H3, a lysine (K for short), abbreviated as H3K9me3, is an epigenetic mark that tells the cell that this nucleosome is part of constitutive heterochromatin [1]. In most organisms this type of heterochromatin includes regions near the telomeres and around the centromeres of the chromosomes. Heterochromatin is rather densely packed, while the more open sections, in which most of the actively transcribed genes are located, belong to what is known as euchromatin.

It has been shown experimentally that during DNA duplication the nucleosomes are randomly distributed between the DNA molecules of the two daughter cells [4]. The missing nucleosomes are then refilled by new nucleosomes. As only the old nucleosomes carry epigenetic marks, this process causes the information to be “diluted” by a factor two. The challenge is now to put the marks back onto the new nucleosomes. As an example, consider the H3K9me3 tag for heterochromatin. The parent cell contains blocks of nucleosomes with H3K9me3 marks alternating with blocks without this tag [5]. In humans, the median length of such a block is about 10 kilobases [6], which corresponds to about 50 nucleosomes. It is not known how the daughter cell puts back the marks onto the heterogenetic blocks in a reliable and robust way.

Models put forward to describe the heterochromatin formation and self-maintenance typically take a one-dimensional view in which heterochromatic marks spread along the nucle-osome chain. At the borders to euchromatin, barrier insulators are postulated [6, 7], which prevent the further spread of the marks. Such insulators contain e.g. DNA sequences that bind proteins, which in turn recruit enzymes that modify nucleosomes such that they can no longer obtain the H3K9me3 mark, see for example [6, 7]. However, this view neglects the fact that chromatin is folded in three dimensions and the enzymes could jump to other sections of the chromosome nearby. Barrier elements alone might thus not be sufficient to contain the spread of heterochromatin. Instead, there must exist some other mechanism that isolates euchromatin from heterochromatin in a three-dimensional context. This mechanism has to overcome various challenges: It needs to provide a physico-chemical environment, a “reaction chamber” in which the corresponding enzymes put back the H3K9me3 marks onto the “right” nucleosomes. There needs to be a physical interface defining a boundary between the nucleosomes that belong to the heterogenetic and the euchromatic domains. Moreover, this physico-chemical environment, the heterochromatin-phase, has to be stable with respect to the reduction/increase of the number of epigenetic marks by a factor of two. Also, the concentration of proteins might fluctuate to some extent during the cell-cycle, thus some stability of this environment with respect to at least minor changes of the protein concentration is required.

What could be a robust mechanism that allows to separate the two types of chromatin and to put the epigenetic marks back to the correct new nucleosomes after duplication? We suggest that such a mechanism requires a particular form of macromolecular condensates, which have meanwhile been observed repeatedly in biological systems [8]. Such droplets, formed by protein and/or RNA molecules, create chemical environments with sharp boundaries against the rest of the cell. It is known for heterochromatin that the protein heterochromatin 1 (HP1) forms droplets at sufficiently high concentrations *in vitro* [9, 10]. These are caused by the attraction between some nonstructured regions in these proteins. In addition, it is known that HP1 binds specifically to the H3K9me3 marks of nucleosomes [11–13]. This suggests that each chromosome with its domains of H3K9me3-nucleosomes and of unmarked nucleosomes acts like a block copolymer at a selective interface between two phases [14]. The blocks containing H3K9me3-nucleosomes would form loops inside the HP1 droplets and the other nucleosomes would loop outside the droplets. Since the boundaries between the two types of nucleosomes would be pinned at the droplet surface, this surface would act as a physical insulator.

If the system is sufficiently robust against changes in parameters, the daughter cells’ chromosomes with their half-diluted marks would still maintain similar configurations regarding loops being in- or outside the droplets after cell division. If the enzyme that puts the marks back to the nucleosomes would only work inside the droplets, the new nucleosomes inside the heterochromatic blocks would recover the proper H3K9me3 marks, while the nucleosomes in the euchromatic loops outside the condensate would remain unaffected. In fact, the H3K9 methylase Suv39h1 is found associated with HP1 [15, 16]. Similar mechanisms may be at work in other droplets, e.g. for the modification H3K27me3 which marks facultative heterochromatin (heterochromatin that is compact only in a subset of cell types). In this case, the role of HP1 might be played by the Polycomb complex PCR1, a protein complex known to have the ability to form condensates and to have specific attraction to H3K27me3 marks [17]. Therefore systems of droplets that organize blocks of nucleosomes into sets of loops might serve as a platform to confer epigenetic memory across cell generations.

The scenario proposed above requires liquid-liquid phase separation and the formation of protein condensates, which has recently attracted intense research in the physics of life. From the point of view of equilibrium phase transitions, two aspects are puzzling: First, why do condensates form from individually water-soluble components? Second, why do condensates have characteristic sizes, i.e. they do not seem to be subject to Ostwald-type ripening? While the first aspect can be explained by attractive interactions between different components [18], in some cases involving mRNA, the second aspect is often considered as intrinsically non-equilibrium in nature. For the case of HP1 domains, both aspects could be explained at the same time if the condensates require methylated chromatin to form. This is consistent, for instance, with the observation that in early *Drosophila* embryos, the total intensity of fluorescently tagged HP1 remains the same over several cell cycles, regardless of the formation of HP1 condensates [10]. This suggests that it is the presence of methylated heterochromatin and not the HP1 concentration alone that controls the formation of HP1 condensates.

In this work we show that binary liquids which are set in a fully miscible state form stable droplets upon weak interactions with long polymers chains, where the size of the condensate is controlled by the size of the collapsed chain or sequences of the chain in equilibrium. We call this mechanism polymer assisted condensation (PAC). We further show that this state is robust with respect to changes of interaction parameters. The latter is important for the function of heterochromatin since after cell-division only half of the methylated nucleosomes are present. Furthermore, our model allows for small changes in the environmental concentration of the condensing protein. We aim to present a generic model using concepts of polymer physics and reveal its properties analytically in its simplest form, solve it numerically in general, and proof the concept by molecular dynamics simulations.

## II. ANALYTICAL MODEL FOR POLYMER-ASSISTED LIQUID-LIQUID PHASE SEPARATION

We start to outline a simple model for the condensation of proteins in the presence of a weakly attractive polymer. Although it can be applied to any two-component solution in interaction with a polymer phase, for simplicity we denote the solvent component which selectively interacts with the polymer by HP, and the common solvent as water. We further assume a binary interaction between HP and water denoted by *χ*. We consider a bulk state outside of the polymer region which consists of HP at concentration (volume fraction) *c_b_* in water which is well above the critical point of demixing, *χ* > *χ_c_*, but in the mixed state outside of the coexistence region, as indicated by the blue circle on the lhs in Fig. 1. This situation is different from that previously considered by Brochard and de Gennes [19], in which the solvent mixture is above the critical point at which fluctuation effects dominate and metastability does not occur. The essential idea is that the preferential interaction between HP and the polymer, which we call *ϵ* in the following, shifts the stability of the solution to the condensed state, which in turn is limited by the resulting coil size of the polymer. Our model assumes thermal equilibrium and no active or intrinsically dissipative processes are necessary.

**Figure 1.**
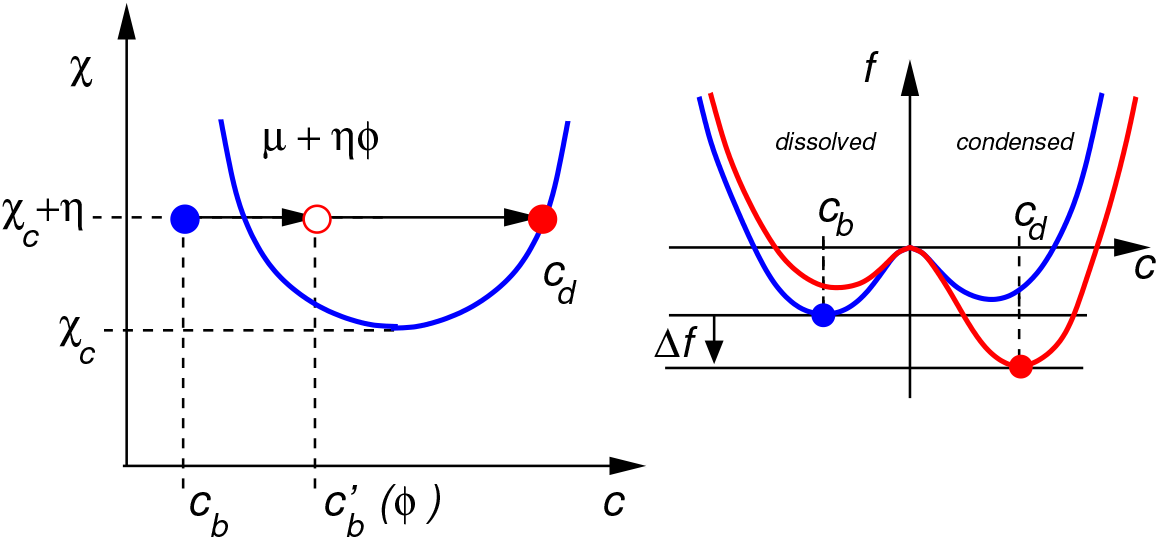
Sketch of the phase diagram of a two-component solution with the HP-concentration given by *c* (lhs). The coexistence states are indicated by the blue line and the critical interaction parameter is given by *χ_c_*. The blue circle denotes the bulk state with concentration *c_b_* outside of the coexistence region. The presence of the polymer with monomer concentration *ϕ* shifts the effective chemical potential to a higher value, which can be associated with a virtual increase of the bulk concentration, and gives rise to phase separation of the solution with the stable phase of an HP-polymer-droplet at concentration *c_d_*, shown by the red circle. The rhs illustrates the free energy profile as a function of the concentration: In the bulk the dissolved state is stable and the condensed state is meta-stable (blue line). The polymer shifts the effective chemical potential (the external field in the context of phase transitions) resulting in a discontinuous transition to the stable condensed state where the dissolved state is meta-stable (red line). The free energy gain, Δ*f*, due to the interaction with the polymer is given by the arrow.

The free energy per volume unit inside the polymer coil in the Flory-Huggins approximation reads:

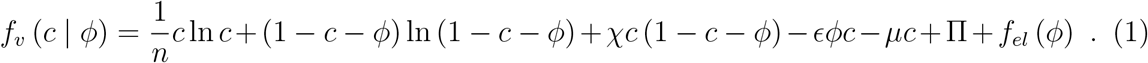

Here, *ϕ* denotes the monomer volume fraction and *c* denotes the volume fraction of HP, *μ* is the chemical potential of the bulk HP, and Π is the osmotic pressure of the bulk phase. Unless otherwise stated, we use *k_B_T* as the unit for the energy, and the volume unit is given by the size of the solvent molecules in the spirit of the Flory-Huggins (FH) lattice model. In general, the size of HP can be *n* times the size of water. Since we are interested in the general physical understanding of the model we restrict ourselves here to the symmetric case, *n* =1, which simplifies the analytical arguments substantially. The term *f_el_*(*ϕ*) denotes the free energy penalty associated with a swelling of the polymer. It will only be of importance for the numerical solution of Eq. (1) and is given by *f_el_*(*ϕ*) = *α*(*Nϕ*^2^)^−1/3^ with a numerical prefactor, *α*, of order unity.

We note the following relations for the bulk state (*ϕ* = 0): 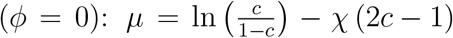, Π = −ln (1 − *c*) − *χc*^2^, and the Gibbs-Duhem relation: Π = *μc* − *f_b_*(*c*), where *f_b_* denotes the free energy per unit volume of the bulk. The symmetry of the bulk solution is broken by *μ* only, thus the coexistence line is defined by *μ* = 0 and given by

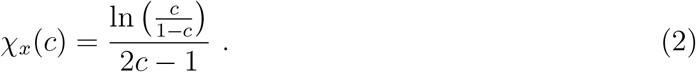

This is sketched in Fig. 1 on the lhs with the blue line. The critical point is given by *c_c_* = 1/2 and *χ_c_* = 2. We introduce the parameter

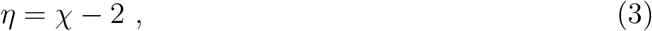

which is taken as positive in the following. A completely dissolved bulk state of HP is given by *μ* < 0 for *η* > 0 which is sketched by the blue line on the rhs of Fig. 1.

We now turn to the free energy inside the polymer volume. With respect to the symmetry of the bulk phase at *c* = 1/2 we introduce the variable *δ* as

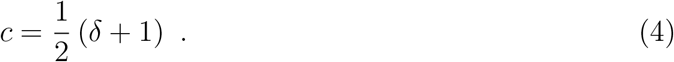

In the spirit of the Landau-model we expand the free energy in Eq. (1) with respect to *δ* and *ϕ* and we sort the terms in the order of the powers of *δ* as follows:

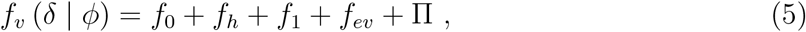

with

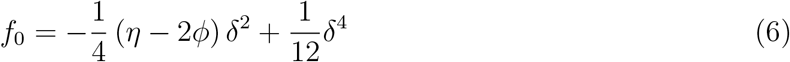

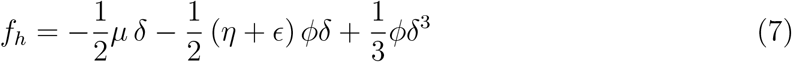

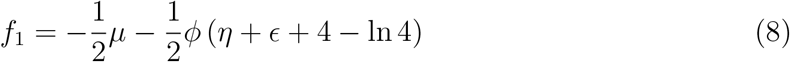

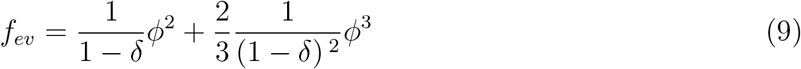

The symmetric contribution *f*_0_ represents the non-trivial bulk free energy at phase coexistence, i.e. in the absence of the field (chemical potential).

We can read-off some interesting physics from the Landau-type expansion. First, without the symmetry-breaking contribution, *f_h_*, we obtain the phase-coexistence located at ±*δ*_0_ with

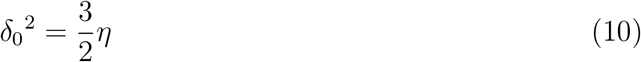

This corresponds to the volume fractions of the diluted (−) and the condensed (+) phases of the bulk at phase coexistence.

The symmetry breaking field is given by *f_h_* = −*hδ* with

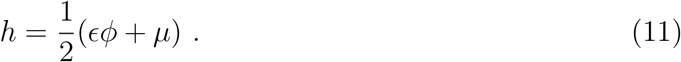

Here, we have used the approximation 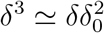 in *f_h_*. The bulk behavior is given by *ϕ* = 0, and *μ* < 0. Thus, for the bulk state we have *h* < 0 and the lower minimum close to −*δ*_0_ is dominating, see blue line on the rhs of Fig. 1. With increasing polymer concentration, the condensed phase becomes the stable one for *ϕ* > *μ/ϵ*. One can regard the polymer field, *ϵϕ*, as shifting the effective chemical potential of the bulk to a virtual concentration 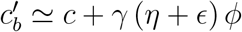 which is located in the coexistence region, as indicated by the red open circle on the lhs of Fig. 1. Here, *γ* denotes a strictly positive prefactor. We note that the polymer field shifts the critical point too, see Eq. (6). However, this effect would be only of interest if *ϕ* ≃ *η*/2. In the following we will consider the case *ϕ* ≪ *η* and disregard this shift.

The terms in *f*_1_ correspond to *δ*-independent contributions. The absolute contribution of the chemical potential due to the shift from the *c*- to the *δ*-variable can be disregarded. The second negative term in *f*_1_ corresponds to the reduction of the mean-field interaction for *δ* = 0. We note that the contribution 4−ln 4 is due to the additional mixing entropy of HP in the binary water-polymer environment. The excluded volume contribution correspond to the usual expressions in the reduced volume for the polymer in the presence of HP, i.e. *ϕ′* = *ϕ*/(1 − *c*).

In order to find the equilibrium state of the polymer-phase with respect to the bulk phase, first we obtain the global minimum of *f_v_* (*δ* | *ϕ*) for a given value of *ϕ*, which we denote as *f_v_*(*ϕ*). The zero-order approach is to the consider the location of the two minima for *h* = 0, given by *δ*_0_, see Eq. (10). Then, the leading order contribution to *f_v_*(*ϕ*) is given by

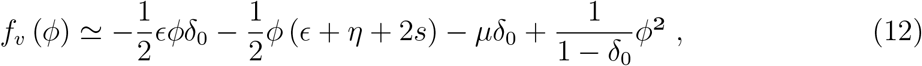

where we have introduced the entropic contribution to the polymer field by *s* = 2−ln 2. We note that we consider *μ* < 0 and thus the bulk contribution has to be taken at *δ_b_* = −*δ*_0_, leading to an additional term 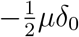. The osmotic pressure, Π, see Eq. (1), corresponds to the negative free energy per volume unit of the bulk at given chemical potential. Thus, *f_v_*(*ϕ*) is nothing but the difference between the polymer free energy and the bulk free energy. Here, we have restricted the excluded volume interaction to the second virial coefficient. We note that at this level we can also take into account the Des Cloizeaux result *f_ev_* = *aϕ*^9/4^ with a numerical constant *a* instead [20].

In order to find the equilibrium polymer volume fraction, *ϕ*, we have to consider the total free energy difference of the droplet with respect to the free energy of the bulk in the same volume. Since the polymer volume is given by *V* = *N*/*ϕ* and thus a function of *ϕ*, the free energy per monomer, i.e. *f*(*ϕ*) = *f_v_*(*ϕ*)/*ϕ* has to be minimized instead. The minimum of the free energy per monomer, *f*(*ϕ*), is located at

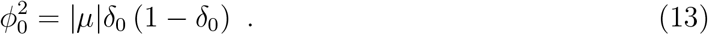

We note that the first two terms in Eq. (12) are proportional to *ϕ*, so they become constant in *f*(*ϕ*) which is why the location of the minimum does not depend on *ϵ*. We note that the value of *ϕ*_0_ is limited from below by the unperturbed coil-size in good solvent: *ϕ*\* ~ *N*^−4/5^, which sets the minimum value |*μ*|* ~ *N*^−8/4^ which is a very small number for large polymers with *N* = *O*(1000). We can check the validity of the assumption *ϕ* ≪ *η* made above which reads: |*μ*| ≪ *η*^3/2^. Thus, for values far away from the coexistence point the shift of the critical point due to the polymer field has to be considered. In the inset of Fig. 2 we display the numerical result for the polymer concentration as a function of the absolute value of the chemical potential of the HP-solution for selected parameters of *ϵ* and for *χ* = 2.2. All numerical results are obtained by finding the global minimum of the free energy per monomer unit derived from Eq. (1), i.e. for *f*(*c* | *ϕ*) = *f_v_*(*c* | *ϕ*) /*ϕ*. A nearly linear behavior between the square of the equilibrium monomer concentration and the chemical potential can be observed for smaller values of −*μ*.

**Figure 2.**
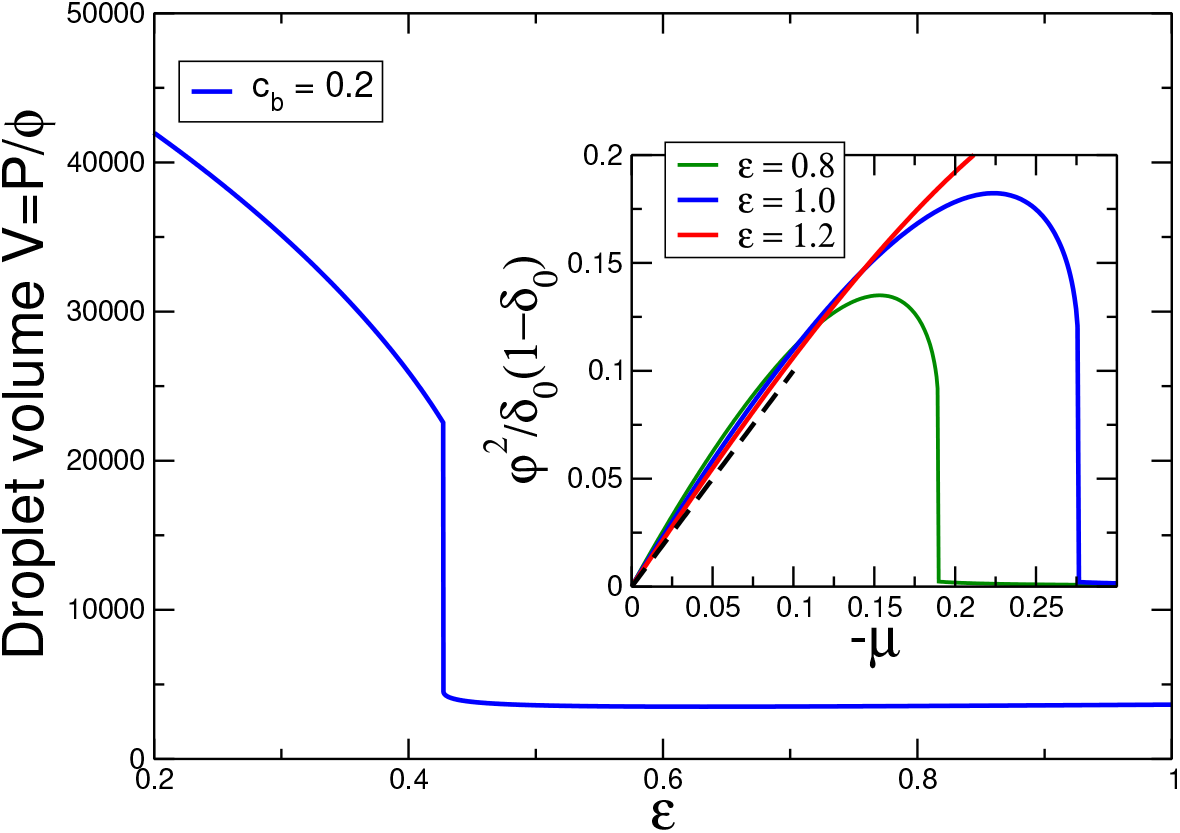
Volume of polymer vs. strength of the HP-polymer interaction. The HP solution is set to *χ* = 2.2 (*η* = 0.2) and *c_b_* = 0.2 well below the bulk condensation point at *c_x_* ≃ 0.25 according to Eq. (2). The degree of polymerization used in the numerical solution is *N* = 500. In the condensed state (*ϵ* > *ϵ_x_* ≃ 0.43) the droplet volume is nearly independent of *ϵ*. Inset: Rescaled squared monomer concentration vs. absolute value of the chemical potential of the HP solution for *ϵ* = 0.8, 1, 1.2. As predicted in the approximate solution, Eq. (13), dashed line, the polymer concentration rises with decreasing HP concentration (increasing absolute value of the chemical potential) and is nearly independent of the interaction with the polymer.

According to Eq. (13) the volume of the condensate droplet, *V_d_* ~ 1/*ϕ*_0_ ~ |*μ*|^−1/2^, shrinks with decreasing HP-concentration in the bulk. That means, that very close to the bulk coexistence, |*μ*| = 0, droplets of maximum size are formed which decrease in size up to the transition, estimated by Eq. (14). While this seems intuitive (smaller droplets for smaller HP-concentration), it means that at the transition (minimum HP volume fraction in the bulk) the maximum degree of collapse of the polymer is observed and that the polymer-droplet expands once again upon adding more HP to the solution.

Given the optimal value of the polymer concentration we localize the transition to the condensed state by using Eq. (11) and *h*(*ϕ*_0_) = 0:

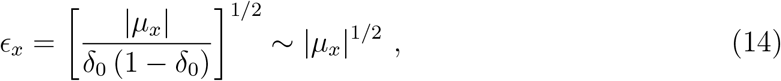

where the index “*x*” indicates the transition between the formation and dissolution of polymer assisted condensates.

A second conclusion taken from Eq. (13) regards the *ϵ*-dependence of the droplet size, which is absent in this approximation. Indeed the variation of the droplet-size with respect to the HP-polymer-selectivity is rather small as displayed in Fig. 2 (behavior right of the transition point at *ϵ_x_* ≃ 0.43). This result is interesting since it shows the robustness of the droplet properties with respect to the interaction between HP and polymer: above a minimal value, which marks the condensation transition, the droplet size only expands weakly with stronger interaction. We note that the fraction of methylated nucleosomes is reduced by a factor of 2 after replication, which should be still sufficient to form the heterochromatin state, which in turn can recruit fresh methylase into the HP-rich droplets.

In the upper part of Fig. 3 we display the relative HP-concentration in the polymer volume for various values of the HP-polymer interaction as obtained from the numerical solution. As expected, below the PAC transition, the ratio is very close to unity since the weak HP-polymer interaction cannot bind any substantial amount of HP without the collective effect of HP. This is followed by a jump-like increase above the transition point. The excess of the HP in the condensate is of the order 2-3 for the parameters chosen here. We note that due to the polymer collapse the total volume fraction of the condensate (HP+polymer) is typically of the order 0.6 − 0.8.

**Figure 3.**
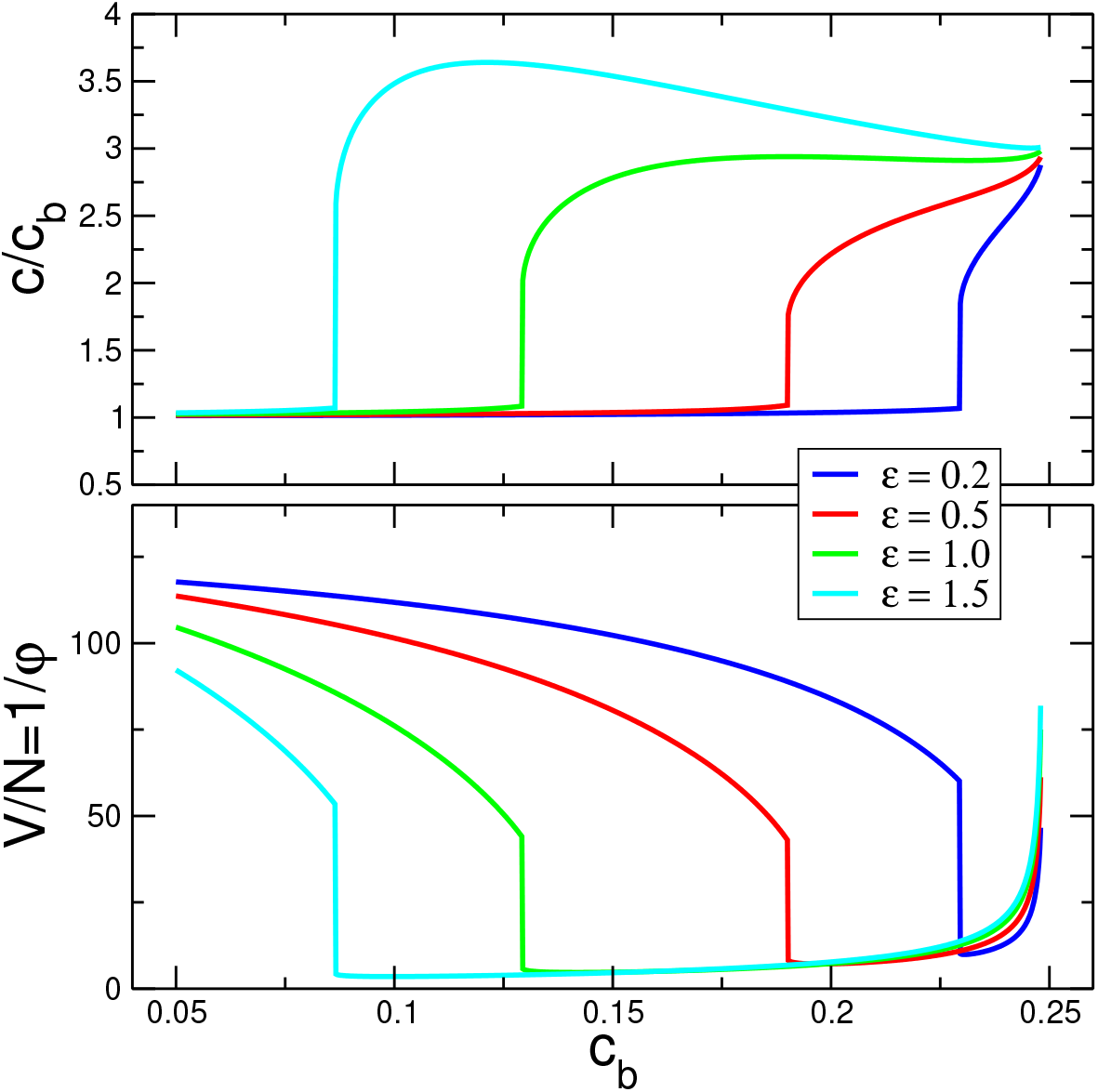
Upper panel: relative HP volume fraction in the polymer volume vs. bulk volume fraction, *c_b_*. Lower panel: Droplet volume per monomer unit vs. *c_b_*. The HP solution is set to *χ* = 2.2 and the bulk condensation point is located at *c_x_* ≃ 0.25

The volume of the polymer coil in monomer units vs. the HP concentration in bulk is displayed in the lower panel of Fig. 3. Increasing the HP concentration to the coexistence condition, see approximate solution in Eq. (14), leads to a jump-like contraction of the polymer along with the formation of the HP-polymer-condensate. Further increase of the amount of HP in the bulk leads to a smooth increase of the droplet volume which is associated with an increase of the HP-fraction. Most significant is the drop of the HP-concentration at the transition point caused by the collapse of the polymer and the relative increase of the polymer volume fraction. This is followed by a steady increase of the HP volume fraction within the droplet which becomes the majority fraction of the co-condensate. The behavior is displayed nearly up to the bulk coexistence point, *c_x_*.

On the lhs of Fig. 4 we show the phase diagram as predicted by Eq. (14). The surface corresponds to the function *ϵ_X_*(*c_b_, χ*). The PAC scenario is bounded by the bulk phase transition for larger values of *c_b_* and *χ* respectively, see Eq. (2), as indicated by the blue line. The rhs of Fig. 4 displays a comparison between the numerical solution of the full free energy model (data points) and the Landau-type approximation (yellow line) for the case *χ* = 2.2. We note that the numerical solution can be easily extended to the case of larger size ratio of HP to water by considering *n* > 1 in Eq. (1).

**Figure 4.**
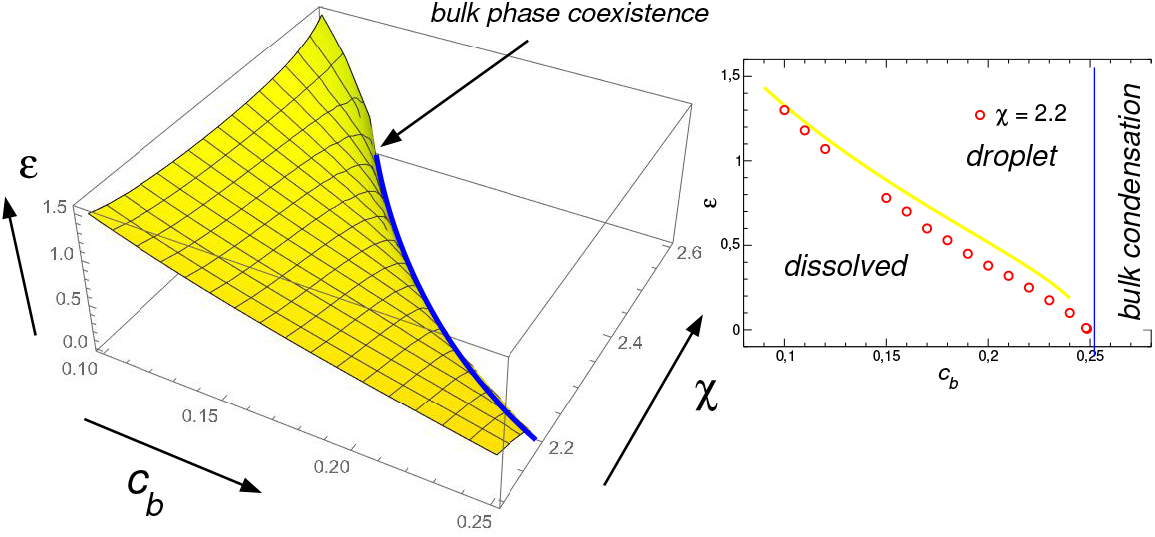
Left: Phase diagram as predicted by Eq. (14). The blue line marks the bulk phase transition in the (*c_b_, χ*)-plane. States above the surface are polymer-assisted condensates. Right: Phase coexistence obtained from the numerical solution for *χ* = 2.2 (circles) compared with the analytical approximation from Eq. (14). The vertical blue line indicates the bulk phase transition.

## III. MOLECULAR DYNAMICS SIMULATIONS

To test the predictions of our mean-field theory, we have carried out molecular dynamics simulations using a standard bead-spring model for the polymer chain, and representing HP molecules by unconnected beads (of the same diameter as the monomers) in the background of an implicit solvent model. The interaction model involves attractive Lennard-Jones (LJ) potentials between the HP-beads, characterized by *χ_S_*, and between the monomers and HP, characterized by *ϵ_S_*. Here, the index “S” indicates the values used in the simulation model. First, we note that due to the hard-core repulsion, a minimum value of *ϵ_S_* is necessary to realize a crossover from repulsion to adsorption of the HP-beads with respect to the polymer chain. This has been studied with the same simulation model in a previous work [21], see Fig. 4 therein, and is given by *ϵ*_*S*0_ ≃ 0.6. The bulk phase diagram of the LJ-system has been studied before [22] and the critical point was found to be located at *χ_X_* ≃ 0.9 and *c_X_* = 0.32. We used a simulation setup with *χ_S_* = 1.1, well above the critical point. The transition to the condensed phase is located at about *c_b_* ≃ 0.0575, while the chain length in our simulations is *N* = 300. Simulations are carried out using the LAMMPS molecular dynamics package [23].

The main results from our simulations are displayed in the phase diagram in Fig. 5. We observe PAC at HP-concentrations well below the bulk phase coexistence, to the left of the vertical red line, as indicated by the open circles. In these states droplets are formed in equilibrium with the surrounding bulk which are restricted in size by the polymer. PAC is bounded by a concentration-dependent HP-polymer interaction, *ϵ_X_*(*c_b_*), which is sketched by the dashed blue line in the phase diagram. As compared to the symmetric FH-model the condensation is shifted to lower values of the HP concentration. This is a consequence of the asymmetry of the LJ-system (spheres vs. implicit solvent). Otherwise, we can recognize all features as predicted from the mean-field model, see Fig. 4. The snapshots display a sharp boundary between the droplet and the bulk phase, and droplets are usually dominated by highly elevated concentrations of HP-molecules. An exception occurs at very low HP-bulk-concentrations at which, close to the phase boundary, the collapsed polymer phase contains only a minority of HP-molecules (red dots in the phase diagram). Here, HP plays the role of a gluonic solvent which forms temporary bridges between monomers as described in Ref. [24]. We note that such a state of the HP-polymer-condensate corresponds to the one assumed in a recent work by Spakowitz and coworkers [25].

**Figure 5.**
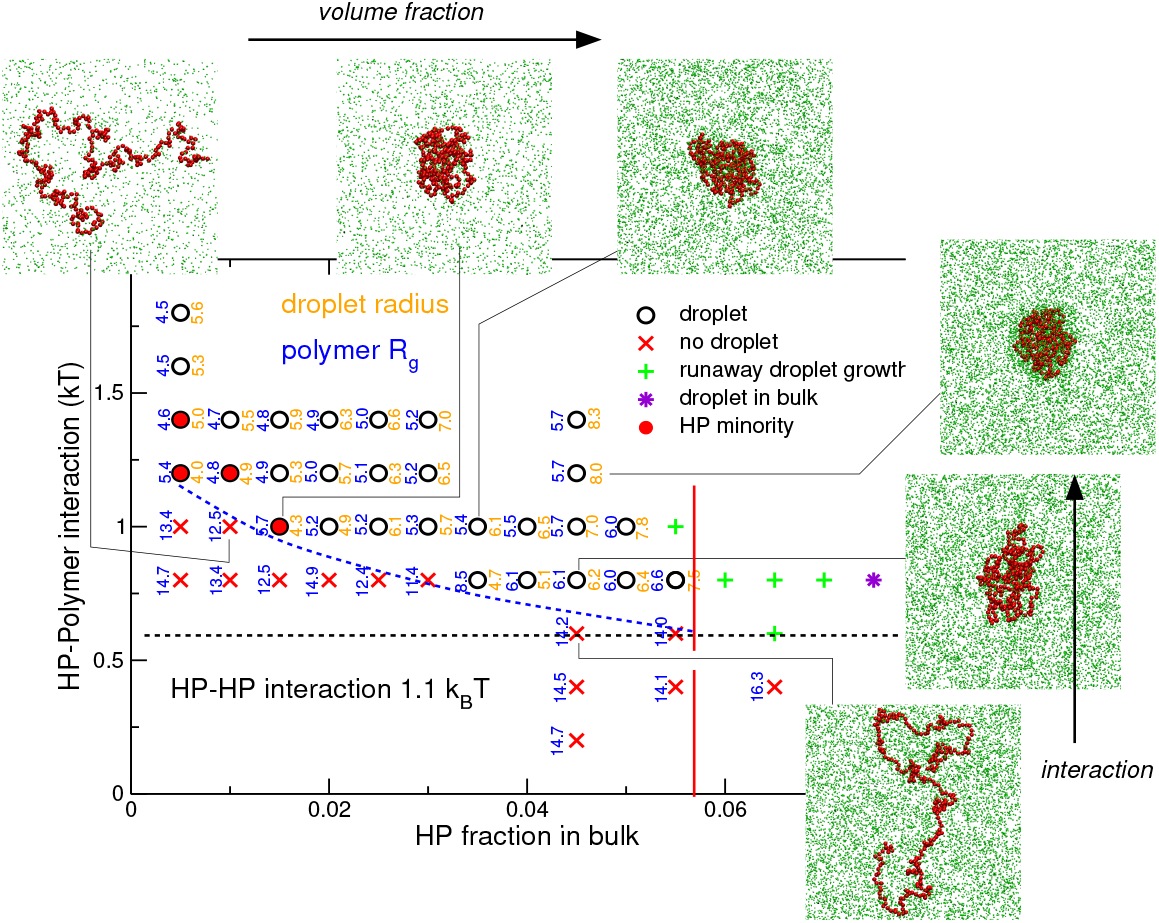
Phase diagram of the simulated system for a polymer chain of length *N* = 300. The LJ-interaction parameter among HP-beads is chosen as *χ_S_* = 1.1, well above the critical point. The bulk condensation transition is indicated by the red vertical line. The blue dashed line sketches the phase boundary between the droplet and the dissolved state, as predicted in Fig. 4. The dashed horizontal line indicates the threshold to HP-polymer adsorption, i.e. *ϵ* = 0 in the mean-field model. Circles indicate droplet states while crosses symbolize dissolved states. Points to the right of the red line belong to bulk condensed states. Snapshots from the simulations are displayed for selected parameters. The numbers on both sides of the data points denote the radius of gyration of the polymer (blue) and the droplet-radius (orange), respectively.

In Fig. 6 we display the droplet volume as defined by 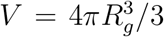, where *R_g_* denotes the radius of gyration of the polymer chain. By crossing over the transition point at about *ϵ_X_* ≃ 0.7, the polymer collapses and the droplet volume turns nearly independent of the HP-polymer interaction as predicted by the mean-field model.

**Figure 6.**
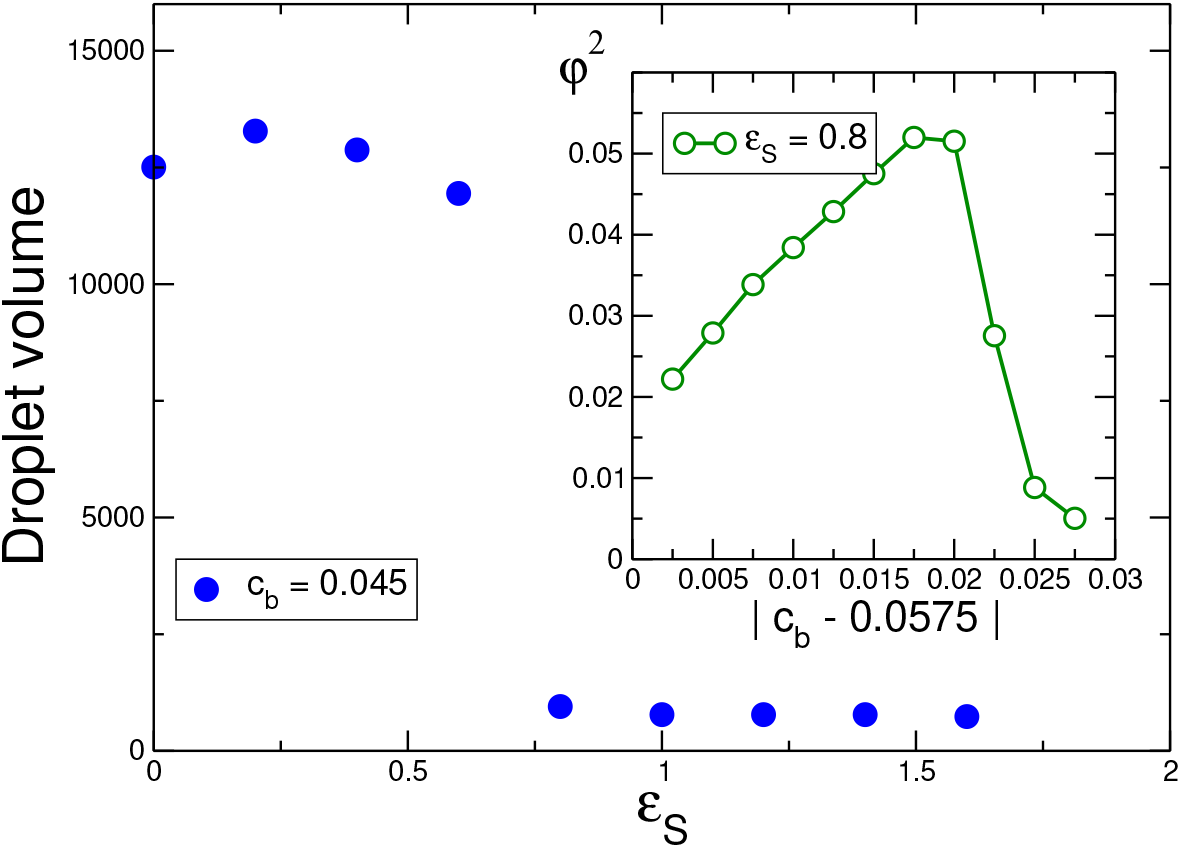
Volume of polymer droplet vs. strength of the HP-polymer interaction in simulation units. The HP bulk solution is set to *c_b_* = 0.045, well below the bulk condensation point at *c_x_* ≃ 0.0575. The degree of polymerization is *N* = 300. In the condensed state (*ϵ* > *ϵ_x_* ≃ 0.7) the droplet volume is nearly independent of *ϵ_S_*. Inset: Squared monomer concentration vs. distance to the condensation point, |*c_b_* − *c_x_*| of the HP solution for *ϵ_S_* = 0.8.

Following the result of the analytical section, see in particular Eq. (13) and the inset in Fig. 2, we have calculated the squared monomer density vs. the distance of the bulk concentration to the coexistence point, Δ*c* =|*c_b_* − *c_x_*|. Here, we have taken the plateau of the monomer concentration profile to calculate *c*. Since we do not have an analytic expression for the chemical potential of the LJ-fluid, we make use of the fact that close to the coexistence point, the chemical potential varies linearly with Δ*c*, i.e. *μ* = *μ_X_* − *a*Δ*c* with some constant *a*, and *μ_X_* denoting the chemical potential at coexistence (which is zero for the symmetric FH-model). The result is displayed in the inset of Fig. 6 and can be compared to the prediction of the theory as shown in the inset of Fig. 2. As predicted, the polymer exhibits its maximally collapsed state at the transition point at the lowest HP volume fraction (larger values of Δ*c*), and then expands when the amount of HP is increased up to the bulk coexistence point (Δ*c* → 0).

To conclude this section, we have shown that PAC of a binary fluid can be observed in molecular dynamics simulations. Although a direct quantitative comparison is not easily possible due of the difference between the equations of state for the LJ-model and the FH-model, all the features predicted by the simple analytical model are displayed semi-quantitatively in the simulated system, including the phase diagram. In particular, the robustness of the properties of the HP-polymer droplets with respect to the HP-polymer interaction is also shown in the simulated model.

## IV. DISCUSSION AND CONCLUSIONS

What is most remarkable about the presented model of PAC is the fact that a discontinuous phase transition is induced in a small system, the extension of which is strictly limited by the polymer size. In fact, the HP-polymer droplet cannot nucleate a macroscopic phase transition in the bulk, since the condensed phase is stable only in the presence of a polymer. Our model can be extended to copolymers where individual sequences are tagged as HP-affine (*ϵ* > 0) while the other monomers behave neutral or repulsive with respect to HP. An example is shown in Fig. 7 for the case of a 9-block copolymer. The only difference being the free energy effort of loop formation which is of order *k_B_T* ln *N_b_* per loop, where *N_b_* denotes the number of monomers per HP-repulsive block. Again, the core of the unimolecular micelle is formed by PAC and contains a major fraction of HP.

**Figure 7.**
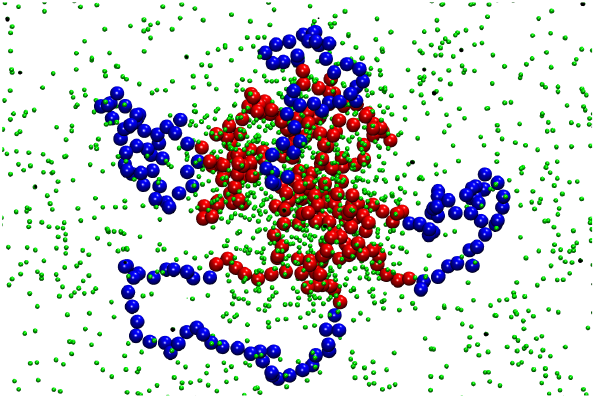
Snapshot of a block-copolymer which is composed of 5 blocks of HP-affine monomers (*ϵ_S_* = 1.0, colored red), and of 4 blocks of HP-repulsive monomers (*ϵ_S_* = 0, colored blue) immersed in a solution with HP of volume fraction *c_b_* = 0.045. Each block contains 40 monomers. The HP-affine blocks form the core of a unimolecular micelle driven by PAC. HP molecules are displayed in green.

Let us note the similarity of the result for the polymer coil size with respect to the HP concentration, lower part of Fig. 3, with the well-known co-nonsolvency effect observed in some polymers [24, 26–29]. Some cosolvents, frequently low alcohols such as methanol and ethanol, when admixed to water can cause a collapse of polymers such as PNiPAm in an intermediate concentration region with a similar re-entrance behavior of the polymer volume as shown here. The difference with respect to PAC is that the alcohol-water mixtures are perfectly miscible (*χ* ≃ 0) and the polymer collapse is a result of an effective monomer-monomer-attraction induced by the non-specific preferential attraction of the cosolvent by the polymer, which can be associated with the formation of temporary co-solvent-bridges between monomers [28, 29]. We note the close similarity with the heterochromatin model proposed in Ref. [25]. This mechanism has been termed as “gluonic” in a previous work [24]. In this case the gluonic solvent (cosolvent) is essentially bound to the monomers and higher values of *ϵ*, typically of order 2 *k_B_T* are necessary [21]. In the case of PAC already very weak preferential interactions down to *ϵ* ≪ 1 *k_B_T* between HP and the polymer can trigger a stable droplet formation. The computer simulations presented in this work give indications for the gluonic regime in the case of very low HP-concentration, see Fig. 5. It is worth noting that both mechanism can work for protein condensates since the difference is only the miscibility of the protein with water and the affinity of the protein with respect to the polymer.

A particularly interesting aspect of PAC is that HP forms the major fraction in the droplet volume which therefore leads to its fluid-like property [30], while the gluonic mechanism leads to a gel-like polymer scaffold which hosts a smaller fraction of the rather strongly bound protein. This is related with a well defined fluid phase boundary of the PAC droplet. Furthermore, the fluid droplet can form a reaction container for other molecules and enzymes which are soluble and even attracted to the HP-enriched phase. Another difference concerns the robustness of the droplet with respect to drastic changes in the polymer-protein interaction, which we have denoted here as *ϵ*. Only low values of *ϵ* < *k_B_T* are necessary for PAC but also larger values result in a liquid protein phase in PAC since the dense phase of protein around the polymer leads to saturation of adsorption and not to strong coupling effects between the monomers. By contrast the gluonic binding requires higher values of *ϵ* to occur and leads to strongly immobilized proteins which form transient bonds between the monomers.

These insights suggest that the PAC scenario provides a good starting point for describing the properties of heterochromatin, particularly how it can be inherited across cell generations. At this point it remains to be seen whether the PAC system with a block polymer alone is sufficient for the reliable inheritance of heterochromatin or whether additional components (e.g. insulators as described in the introduction) need to be present to prohibit an increasing number of euchromatic nucleosomes getting sucked into the droplets over time to become enzymatically tagged as heterochromatic. We plan to address these questions in a future study.

It is useful to discuss the model by Spakowitz and coworkers for epigenetic inheritance in some detail [25]. The model was originally developed to calculate the contact map of an entire human genome [31]. Such contact maps are now routinely measured by chromosome conformation capture techniques [32]. An experimentally determined methylation profile served as input for a coarse-grained chromatin model with nucleosome resolution [25, 31]. The model explicitly takes into account the HP1 molecules, which bind preferentially to methylated nucleosomes and then form bridges to other bound HP1s nearby. The Monte Carlo (MC) simulations have indeed successfully reproduced an experimental contact map [31]. In the later study [25] this model was then used to investigate the inheritance of epigenetic tags through mitosis. An MC simulation was performed to determine 26 equilibrium polymer configurations and HP1 binding profiles, which were used as an input to calculate steady state methylation probabilities. For the latter, it was assumed that the on-rate of methylation for a nucleosome is proportional to the number of HP1-bound tails near that nucleosomes (while the off-rate is not). Each cycle was considered to be one cell generation. It was shown that the methylation profiles could be inherited stably over various cell generations. To achieve this, however, the HP1 concentration had to be carefully adjusted to the given reaction rates. If the HP1 concentration was changed slightly to larger or smaller values, the proportion of methylated nucleosomes would either increase or decrease with each subsequent cell generation (see Fig. 2 in Ref. [25]). It should be further noted that the chromosome conformations were frozen during the methylation reaction and were determined from the methylation state of the previous cycle, i.e. before cell division, instead of being sampled afresh beginning with a semi-methylated state. This could affect both the conformations and the resulting methylation profiles.

This suggests to us that this scenario does not give the full picture. On one hand, it shows strong bridging interactions between nucleosomes, which would rather lead to a gel-like state that could hinder the free diffusion of e.g. the methylases through the heterochromatic region. On the other hand, a mechanism would be needed that ensures that the right concentration of HP1 molecules is always present to avoid a runaway mechanism. In contrast, the PAC scenario offers liquid-like heterochromatin compartments that are robust against changes in system parameters such as the concentration of HP1 proteins.

Of interest will be to study the methylation dynamics in the PAC scenario across the cell generations. The individual recovery kinetics after dilution of the tags could also show an interesting dynamics in which the radial density distributions within a droplet of the methylated nucleosomes and of the methylases could change over time due to methylation reactions and subsequent changes in chromosome conformation. It is also of interest to find out whether the layer of euchromatic loops at the outside of the HP1 droplets kinetically protects smaller droplets from merging into larger droplets. Finally, it will be very interesting to compare similarities and differences between the mechanisms of maintenance and/or establishment of constitutive and facultative heterochromatin.

## ACKNOWLEDGMENTS

JUS acknowledges support from the DFG under the grant number SO-277/17. HS and JUS was supported by the Deutsche Forschungsgemeinschaft (DFG, German Research Foundation) under Germany’s Excellence Strategy – EXC 2068 – 390729961– Cluster of Excellence Physics of Life of TU Dresden.

## Notes

### Competing Interest Statement

The authors have declared no competing interest.

